# Prescriptions For The Control Of A Clonal Invasive Species Using Demographic Models

**DOI:** 10.1101/2021.12.28.474361

**Authors:** Gabriel Arroyo-Cosultchi, Jordan Golubov, Jonathan V. Solórzano, Maria C. Mandujano

**Author notes:** Correspondence; (J.G.). These authors contributed equally to this work.

## Abstract

Until recently, little focus has been given to determine the population dynamics of invasive species and evaluate their genetic variation. Consequently, not much is known of what drives clonal invasive species and their demography. Here we describe the population dynamics of *Kalanchoe delagoensis* (Crassulaceae), considered invasive to several countries. We quantified the demography of a population in central Mexico using integral projection models (IPM) in a population that reproduced asexually exclusively through plantlets. The effect of clonal recruitment on *λ* was evaluated by changing plantlet survival and simulating management scenarios that used previous data of watering and seven experimental herbicide treatments. The finite rate of population increase indicated that this *Kalanchoe delagoensis* population is growing (above one) with significant potential increases that correlated with water availability. The IPM showed that plantlet survival and recruitment were the most critical steps in the cycle for the population and simulations of different management scenarios showed that reducing plantlet survival significantly decreased *λ* only in two out of the seven herbicides used.

## 1. Introduction

Interest in alien invasive species (AIS) has grown in recent years because the impacts are widely recognized [1], and global costs are being quantified [2–4]. Critical stages of invasion involve the establishment and spread into new habitats (categories C, D, and E in [5]), which can depend on a suite of factors including habitat suitability [6], components of propagule pressure (frequency and size of introductions [7,8]), presence/absence of pollinators, dispersers or herbivores [9] and phenotypic plasticity [10,11]. The reproductive mechanisms behind the spread can vary widely, however, even though clonality seems to be quite common among alien invasive species [12], the role played by clonality during spread is context-dependent [13,14]. To our knowledge, of the 31 papers that deal with population dynamics of invasive species, six had some form of vegetative reproduction [15,16]. Even though no attempt has been made to evaluate the overall demographic importance of clonality among invasive species, it is widely recognized as an essential trait in risk assessments [17–19], invasion success [14], and is a critical component of population growth [20].

Only recently have demographic models been used to assess components of invasion and to guide management of invasive species [15,21–25] which can even include prioritization based upon cost [26]. Population dynamics of AIS commonly have high population growth rates [15], and transient dynamics seem to play a part in invasion success [16,27,28]. Invasive species have also been shown to have demographic plasticity [29,30] and buffer environmental variation [21,31] which enables them to exploit both ends of the phenotypic plasticity spectrum [10, master of one, jack of all] and even combinations of this [32]. Targeting certain life stages (or combinations thereof) can provide insight towards identifying critical pathways [22,25,33,34] that would lead to optimal management strategies [15,27], as well as diminish the cost and overall impact of AIS [35].

Integral projection models (IPM’s) have been used recently and widely for demographic analyses (e.g. [36,37]), including assessments of population viability for endangered plants (e.g. [38]) or invasive plants (e.g. [24,25,31]). Demographic modelling can then provide insight into short-term possible directions for management, and benefit from being robust to small data sets compared to matrix population models ([39] and are able to incorporate other explanatory drivers as covariates [31,38,40].

The genus *Kalanchoe* is native to the dry regions of Madagascar and Eastern Africa, and several species are now found in several countries (Mexico, Australia [listed under the noxious weed act], USA, Spain, and South Africa) where it has been introduced and spread either accidentally or through the horticultural trade [41,42]. In Mexico, several species and one hybrid have been introduced (*K. pinnata*, *K. tubiflora*, *K. blossfeldiana*, *K. fedtschenkoi*, *K. daigremontiana*, *K. calcynum*, *K. laciniata*, *K. integra* and *K. daigremontiana × delagoensis*, [43]); one, *Kalanchoe delagoensis*, can now be found in six states within the arid environments in Mexico and has been declared invasive [44]. *Kalanchoe delagoensis* can hybridize with *K. daigremontiana* to generate what is known as Houghton’s hybrid, also introduced with no evidence of hybridization in Mexican populations even though they may grow sympatrically in some areas [45]. In addition, populations of *K. delagoensis* share a single genetic makeup suggesting a single introduction event with subsequent spread. This means that *K. delagoensis* could establish founder populations under a wide range of conditions entirely through clonality [45]. Negative impacts of *K. delagoensis* include allelopathic effects [41], harm to domestic animals [46–48] and changes in available soil carbon [34]. Even though herbicidal control has been tried, as have controlled burns, management has been difficult due to the high production of clonal plantlets from the margin of leaves. This apomictic trait is the main driver of population dynamics of its congener *K. daigremontiana* in Venezuela [49].

The aims of this study were to 1) assess the population dynamics of *K. delagoensis*, and 2) determine the management effectiveness of targeting the plantlet life stage which has been previously shown to be susceptible to some herbicides.

## 2. Results

### 2.1. Vital rates

A total of 497 individuals were followed during the study period. Models adjusted for survival, fecundity and growth to generate the kernel used a log transformation of plant size to improve fit. Of the models fitted to the data (survival with 5 models, growth 6 models, reproductive probability 8 models and fecundity 8 models), those with the lowest value of AIC were used (Figure 1). The production of plantlets was found for even small individuals that started to produce plantlets within the year and with plant size as small as 1.5 cm. The population of *K. delagoensis* studied showed no seeds and therefore the driver of population growth is entirely through the proliferous production of plantlets. During the period (2010-2011), the survival probability of *K. delagoensis* and growth (regression with slope > 1; Figure 1 a and b) increased with stem size, and probability of reproduction and fecundity also increased with stem size (Figure 1 c and d). In our study site and period, the population growth rate (*λ*) of *K. delagoensis* with plantlet survival set for the control treatment (no water and full sunlight) was > 1 (Table 1), which would suggest the growth rate under severe natural conditions. Fecundity (Fmatrix) contributed 60% and 52% to the population growth rate (*λ*) for 100% water and other herbicides treatment, respectively. The high values of *λ* (3.51 and 2.08) of these models suggest the importance of vegetative reproduction in the dynamics of the population and the potential for invasion. Survival-growth (Pmatrix) contributed 100% and 78% towards *λ* for G/2-4D and 2-4D treatments, respectively basically due to the high plantlet mortality caused by the herbicide treatments. Herbicide treatments (G/2-4D and 2-4D) contributed to the reduction of *λ* and point towards an effective control of *K. delagoensis* populations that rely on clonal reproduction. Although not considered here, the effects of plantlet mortality could be offset by high water availability which suggests herbicide treatments should be avoided during rainy seasons to maximize plantlet mortality.

**Figure 1.**
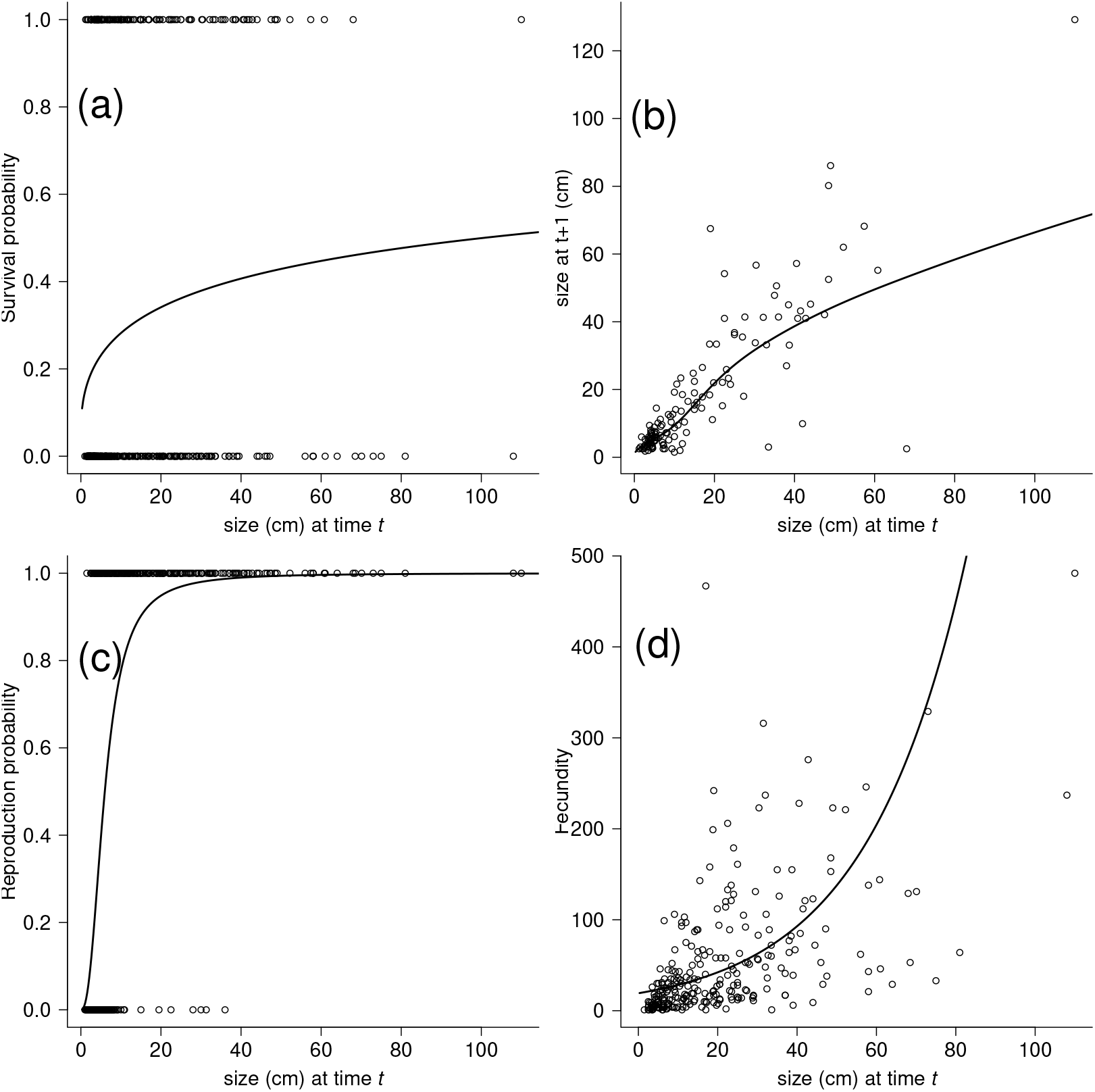
Fitting of the survival, growth, reproduction probability and fecundity of the *K. delagoensis* 2011 data. **(a)**. The survival (s) data are plotted (0, death; 1, survival) as function of individual size *x* (plant height in cm), along with a logistic regression fitted to the data. The fitted curve is log(s/(1-s)= −1.904 + log(0.206)*x* (*P*<0.05). **(b)** The data on year-to-year changes in size, along the regression fit for mean size at year *t*+1 as a function of size in year *t*. The fitted line have *μ*= 0.327+log(0.885)*x* (*P*<0.05) and *σ*^2^= −1.273+log(0.269)*x* (*P*<0.05). **(c)** The reproductive (r) probability are plotted (0, non reproductive; 1, reproductive) as function of individual size *x* (plant height in cm), along with a logistic regression fitted to the data. The fitted curve is log(r/(1-r)= −4.316 +log(2.414)*x* (*P*<0.05). **(d)** The fecundity as a function of individual size, a long with the regression for the mean number of plantlets. The fitted line is *μ* = 2.955+log(0.039)*x* (*P*<0.05). The *y*-axis scales are different among the panels.

**Table 1:**
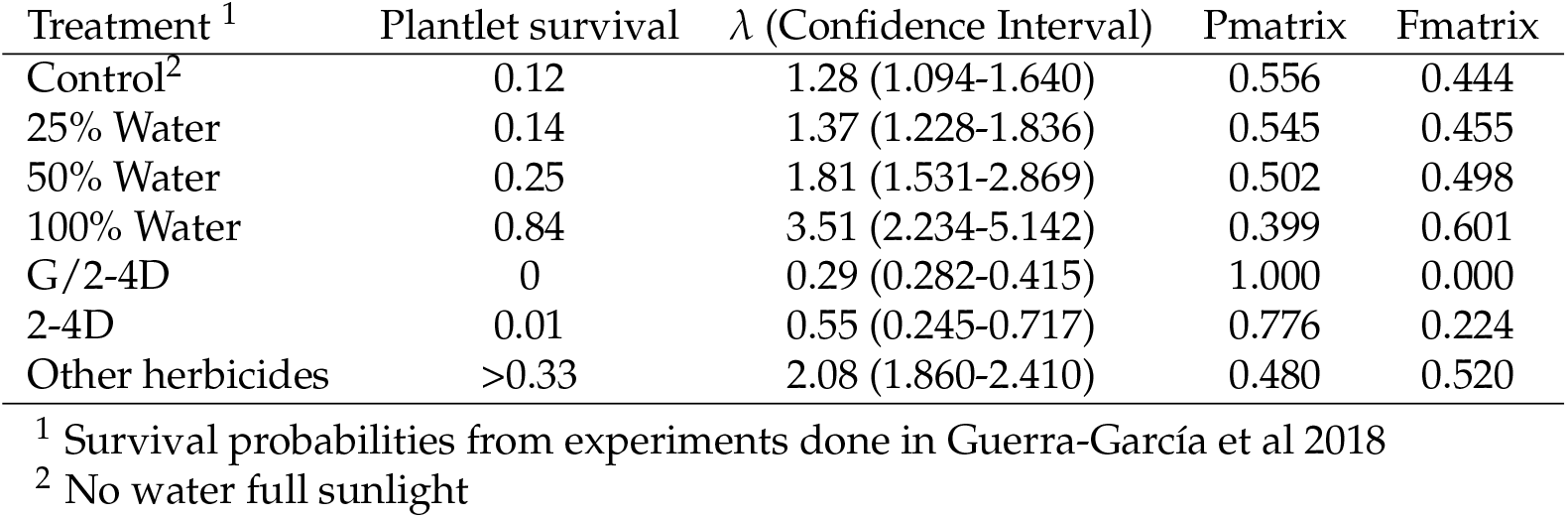
Probability of survival after water and herbicide treatments were applied and the resulting population growth rate with their confidence interval (95%) and the sub-kernels contributions.

## 3. Discussion

Demographic variation at an individual level has only recently been taken into account to determine how individual traits influence population growth [22,50]. Clonality is one of those components that for invasive species seems to be a known and important driver of invasion [14,34,51–53]. However, little attention has been given to the demography of clonal invasive species, and the role played by the clonal component, even though it may be the only means of population growth for some extremely clonal species. This lack of information could be partly due to the cost and difficulty of differentiating genotypes in natural populations of clonal species or to the often overriding importance and haste of direct management options and/or eradication. Nevertheless, among alien invasive species, some examples exemplify the role played by clonality [45,52,54]. Fortunately, sexual reproduction has not been found in the studied population of *K. delagoensis* which the genetic data also suggest [45]. Adding seeds would increase genetic diversity and quite possibly generate even broader responses to environmental variation and increased invasive capacities [54]. Sexual reproduction would also very likely create seed banks such as those found for *K. daigremontiana* [55] complicating management further and increasing management costs basically because of the longer control periods needed to reduce the seed bank. Clonality may not have the potential for dormant stages nor the benefit of genetic variability, but clonal offspring can have higher survival rates over long periods and be a strategy to occupy suitable habitats in short time periods.

Almost all of the invasive species with some form of clonality have population growth rates above equilibrium [15,56–59]. This observation is consistent with the spread of invasive species [5] and is one of the reasons for highlighting the importance of clonality in risk assessments. Clonality coupled with other traits such as self-compatibility [60], resprouting [22], and no need for pollinators or dispersers makes these species more likely to invade successfully than others. There are clear benefits of clonality including the spread of an already successful genotype for a determined habitat. If the traits of clonal offspring have phenotypic plasticity, the species can exhibit a “Jack and Master” strategy [10] allowing the possibility of increasing fitness in favorable environments while maintaining fitness in stressful environments. This would also compound with complex transient dynamics [16,28] and variation in vital rates [31]. Even small increases in favorable abiotic conditions has the ability to add to plantlet survival and therefore increase the invasive potential significantly.

### Management scenarios

A successful management option suggested by [15] was the reduction in fecundity for short-lived invasive species, but seemingly that cannot be applied consistently across species [61]. Reducing one of two demographic processes (fecundity or stasis) was shown to be sufficient to control growing populations [15]. For the sister species *K. daigremontiana* and *K. pinnata* clonality, as well as sexual reproduction, contributed significantly to population growth [34,62]; however, the genetic identity of the individuals was not evaluated. The lack of a sexual component of fecundity in *K. delagoensis* at the site makes control easier and also highlights the danger of triggering higher growth rates if genetically dissimilar individuals are introduced. [15] also found that reducing one demographic process (namely growth or reproduction) under simulated conditions was adequate to control short-lived invasive species, while reduction of at least two was needed to control long-lived perennials. [61] simulated reduction in the sexual component of perennial invasive species and found that reducing this demographic process was not enough to curb population growth. As the plantlets of *K. delagoensis* contribute to fecundity a management scenario would necessarily involve lowering the recruitment of plantlets by using one of two herbicides that had important effects on plantlet survival. Unfortunately not all herbicides had the same effectiveness and of the seven that were tested only two induced high enough mortality in plantlets so generate falls in *λ* needed for adequate control. Lowering recruitment of plantlets, however, poses a problem because plantlets are produced by very young individuals and are generated year-round, such that management must be carried out continuously and would entail long-term control. The timing of herbicide applications should also consider that mortality can be offset in the seasons with higher water availability.

The populations of *K. delagoensis* in Mexico pose an interesting system in which plantlets are the drivers of population growth. This also means that management can be feasible if plantlet survival is severely limited in this population. Little is known of the demographic behavior in other populations which could potentially behave very differently and increase the invasion potential of this and the sister species. Currently, only *Kalanchoe pinnata* and *K. delagoensis* are listed as invasive species in Mexico [44], bit the sister species *K. daigremontiana* and what is known as Houghton’s hybrid (*K. diagremontiana K. delagoensis*) already have large populations with wide distributions and also pose a significant risk as they have very similar demographic behavior.

## 4. Materials and Methods

### 4.1. Species

*Kalanchoe delagoensis* Ecklon & Zeyher [63] is a succulent able to reproduce sexually as well as clonally through pseudobulbils (hence the name “mother of millions”) that arise from the margin of their leaves [64]. Clonal plantlets have accelerated growth, high survival, and a common sight in invaded areas [65]. *K. delagoensis* is considered to be an aggressive invader in Australia and the US [41]. It is included in the exotic species lists in Mexico, and risk assessments suggest a significant risk to native vegetation and soils (Guerrero-Eloisa, unpub. data).

### 4.2. Population Dynamics

For two years (2010-2011), a populations of *K. delagoensis* (hereafter Cadereyta 20.687 061° N *−*99.805 255° W was followed. The site has 15.9°C average annual temperature and 488 mm annual rainfall (data from 60 years obtained from the National Meteorological Service stations at the sites). Vegetation at the site consisted of Chihuahuan xerophytic scrub. During April 2010, 1 × 1 m plots (Cadereyta 7 plots) were set, and all individuals of *K. delagoensis* within the plots were tagged and measured (plant width (cm) and height (cm)). Tagged individuals were followed every two months for a year.

### 4.3. Population growth rate

Demographic data on *K. delagoensis* were obtained as plantlets and adults. Data were widely examined for errors and outliers. All data were used to construct informative models for vital rates (individual survival, growth and fecundity) based on plant size (height, cm). We performed model selection using AICc criteria to choose the most plausible models among a range of different models. The vital rate models support the basis for building of integral projection models [36,37,66] to model the life cycle of *K. delagoensis* ([67]). We used an integrative measure (height, cm) as the size variable and analyzed models assessing the effect of size on each vital rate using generalized linear and Generalised additive models for location scale and shape (GLM, GAMLSS).

The IPM incorporates models for individual survival, changes in size, and fecundity. The models for individual survival used a logit link function GLM and binomial distribution using the lme4 package [68]. We modelled plant growth for year *t*+1 (2011) in relation to plant size from year *t* (2011) with nonparametric normal models using the gamlSS package [69]. We estimated size-dependent fecundity of reproductive individuals as the product of size-specific probability of successfully producing plantlets (estimated with a logit link GLM and binomial errors using the lme4 package [68]), and the size specific number of plantlets produced (with nonparametric normal model using the gamlSS package [69]). This gives an estimate of number of plantlets produced by plants of different sizes as a means of reproduction. Mesh size was set to 500 bins. We numerically integrated the demographic kernel using the mid- point rule to generate the IPM [36]. The dominant with eigenvalue of the square matrix coincides to population growth rate (*λ*). Population growth rates = 1 show population stability, < 1 indicate a population expected to decline and rates > 1 indicate a growing population and over the long term. Confidence intervals (95%) for *λ* were obtained for each population by bootstrapping, such that individuals were resampled to generate 1,000 parameters or each element in the kernel [70].

Possible management scenarios were simulated by changing the survival rates of plantlets. Plantlet survival under different treatments with watering levels and the use of herbicides was used from data taken from [71]. The baseline plantlet survival was the survival of plantlets with no watering and under full sunlight, similar to what would be expected under natural conditions. To assess the contributions of changes in vital rates following management scenarios to *λ* we modified the probability of plantlet survival to what was found in the herbicide treatment experiments. We modelled 3 water treatment (25, 50 and 100% water field capacity) to determine the effect of water on population growth rates and seven herbicide treatments, five of which were grouped into those with more that 33% plantlet survival and those that had the highest mortality (G/2-4 Glyphosate + 2-4D amine mixture and 2-4D, see concentration details in [71]). Population growth rates and error intervals (through boostrapping) were calculated for each treatment. We calculated the population growth rate and the vital rate elasticity and sensitivity for the additive sub-matrices of survival-growth (P) and fecundity (F) [72] based on each IPM representing a different management scenarios (elasticity and sensitivity ∷popbio; [73]).

## Author Contributions

“Conceptualization, JS, GAC and JG; methodology, JS, GAC, JG; formal analysis, GAC; investigation MCM; resources, MCM, JG; data curation JS and JG; writing—original draft preparation JG; writing—review and editing, JG, MCM and GAC; project administration, JG and MCM; funding acquisition, JG and MCM. All authors have read and agreed to the published version of the manuscript.”

## Institutional Review Board Statement

Not applicable.

## Informed Consent Statement

Not applicable.

## Data Availability Statement

The data presented in this study are available upon request from the first author.

## Acknowledgments

We want to thank E. Sánchez and personnel of the Botanical Garden in Cadereyta for providing logistic support. H Altamirano, J Noguez Lugo, H Salgado Garrido and ME Medel Ávila helped during field work.

## References

1. Genovesi, H.; Carboneras, C.; Vila, M.; Walton, P. EU adopts innovative legislation on invasive species: a step towards a global response to biological invasions. Biological Invasions 2015, 117, 1307–1311.

2. Pimentel, D.; Zuniga, R.; Morrison, D. Update on the environmental and economic costs associated with aline-invasive species in the United States. Ecological Economics 2005, 52, 273–288.

3. Bradshaw, C.; Leroy, B.; Bellard, C.; Roiz, D.; Albert, C.; Fournier, A.; Babet-Massin, M.; Salles, J.; Simard, F.; Courchamp, F. Massive yet grossly underestimated global costs of invasive insects. Nature Communications 2016, 7, 12986.

4. Hoffmann, B.D.; Broadhurst, L.M. The economic cost of managing invasive species in Australia. NeoBiota 2016, 31, –18.

5. Blackburn, T.M.; Pŷsek, P.; Bacher, S.; Carlton, J.T.; Duncan, R.P.; Jarosik, V.; Wilson, J.R.U.; Richardson, D.M. A unified framework for biological invasions. Trends In Ecology and Evolution 2011, 26, 333–339.

6. Evangelista, P.H.; Kumar, S.; Stohlgren, T.J.; Jarnevich, C.S.; Crall, A.W.; Norman, J.B.; Barnett, D.T. Modelling invasion for a habitat generalist and a specialist plant species. Diversity and Distributions 2008, 14, 808–817.

7. Duncan, R.P.; Rossinelli, T.M.B.S.; Bacher, S. Quantifying invasion risk: the relationship between establishment probability and founding population size. Methods in Ecology and Evolution 2014, 5, 1255–1263.

8. Lockwood, J.L.; Cassey, P.; Blackburn, T. The role of propagule pressure in explaining species invasions. Trends in Ecology and Evolution 2005, 20, 223–228.

9. Richardson, D.M.; Allsop, N.; D’Antonio, C.M.; Milton, S.J.; Rejmanek, M. Plant invasions–the role of mutualisms. Biological Review 2000, 75, 65–93.

10. Richards, C.; Bossdorf, O.; Muth, N.; Gurevitch, J.; Pigliucci, M. Jack of all trades, master of some? On the role of phenotypic plasticity in plant invasions. Ecology Letters 2006, 9, 981–993.

11. Daehler, C.C. Performance comparison of co-occurring native and aliens invasive plants: Implications for conservation and restoration. Annual Review of Ecology and Systematics 2003, 34, 183–211.

12. Liu, J.; Dong, M.; Miao, S.L.; Li, Z.Y.; Song, M.H.; Wang, R.Q. Invasive alien plants in China: role of clonality and geographical origin. Biological invasions 2006, 8, 1461–1470.

13. Pŷsek, P. Clonality and plant invasions: can a trait make a difference? In The ecology and evolution of clonal plants; de Kroon H J van Groenendael, H., Ed.; Backhuys Publishers: Leiden, 1997; pp. 405–427.

14. Lloret, F.; Médial, F.; Brundu, G.; Camarada, J.; Moragues, E.; Rita, J.; Lambdon, P.; Hulme, P. Specie attributes and invasion success by alien plants on Mediterranean islands. Journal of Ecology 2005, 93, 512–520.

15. Ramula, A.; Knight, T.M.; Burns, J.H.; Buckley, Y.M. General guidelines for invasive plant management based in comparative demography of invasive and native plant populations. Journal of Applied Ecology 2008, 45, 1124–1133.

16. Jelbert, K.; Buss, D.; McDonald, J.; Townley, S.; Franco, M.; Stott, I.; Jones, O.; Salguero-Gómez, R.; Buckley, Y.; Knight, T.; Silk, M.; Sargent, F.; Rolph, S.; Wilson, P.; Hodgson, D. Demographic amplification is a predictor of invasiveness among plants. Nature Communications 2019, 10, 5602.

17. Daehler, C.C.; Carino, D.A. Predicting invasive plants: prospects for a general screening system based on current regional models. Biological Invasions 2000, 2, 93–102.

18. Pysek, M.K.P. Predicting invasions by woody species in a temperate zone: a test of three risk assessment schemes in the Czech Republic (Central Europe). Diversity and Distributions 2006, 12, 319–327.

19. Koop, A.L.; Fowler, L.; Newton, L.P.; Caton, B.P. Development and validation of a weed screening tool for the United States. Biological Invasions 2012, 14, 273–294.

20. Suehls, C.M.; Affra, L.; Médail, F. Invasion dynamics of two *Carpobrotus* (Aizoaceae) taxa on a Mediterranean island: I. Genetic diversity and introgression. Heredity 2004, 93, 31–40.

21. Li, S.L.; Ramula, S. Demographic strategies of plant invaders in temporally varying environments. Population Ecology 2015, 57, 373–380.

22. Sebert-Cuvillier, E.; Paccaut, F.; Chabrerie, O.; Endels, P.; Goubet, O.; Cecocq, G. Local population dynamics of invasive tree species with a complex life history cycle: a stochastic matrix model. Ecological Modelling 2007, 201, 127–143.

23. Williams, J.L.; Auge, H.; Maron, J.L. Testing hypotheses for exotic plant success: parallel experiments in the native and introduced ranges. Ecology 2010, 91, 1355–1366.

24. Ramula, S. Annual mowing has the potential to reduce the invasion of herbaceous *Lupinus polyphyllus*. Biological Invasions 2020, 22, 3163–3173.

25. Zucaratto, R.; Pires, A.S.; Bergallo, H.G.; de Cássia Quitete Portela, R. Felling the giants: integral projection models indicate adult management to control an exotic invasive palm. Plant Ecology 2021, 222, 93–105.

26. Kerr, N.Z.; Baxter, P.W.J.; Salguero-Gómez, R.; Wardle, G.M.; Buckley, Y.M. Ecological predictions and risk assessment for alien fishes in North America. Journal of Applied Ecology 2016, 53, 305–316.

27. Iles, D.T.; Salguero-Gomez, R.; Adler, P.; Koons, P.B. Linking transient dynamics and life history in biological invasion success. Journal of Ecology 2016, 104, 399–408.

28. Horvitz, C.C.; Denslow, J.S.; Johnson, T.; Gaoue, O.; Uowolo, A. Unexplained variability among spatial replicates in transient elasticity: implications for evolutionary ecology and management of invasive species. Population Ecology 2018, 60, 61–75.

29. Claridge, K.; Franklin, S. Compensation and plasticity in an invasive plant species. Biological Invasions 2002, 4, 339–347.

30. Aillie, R.; Reshi, Z.; Wafai, B. Demographic plasticity in relation to growth and resource allocation pattern in *Anthemis cotula*-an alien invasive species in Kashmir Himalaya, India. Applied and Environmental Research 2005, 4, 63–74.

31. Ramula, S. Linking vital rates to invasiveness of a perennial herb. Oecologia 2014, 174, 1255–1264.

32. Sultan, S. Phenotypic plasticity for fitness components in *Polygonum* spcecies of constrasting ecological breadth. Ecology 2001, 82, 328–343.

33. Shea, K.; Kelly, D. Estimating biocontrol agent impact with matrix models: Carduus nutans in New Zealand. Ecological Applications 1998, 8, 824–832.

34. Herrera, I.; Chacón, N.; Flores, S.; Benzo, D.; Marínez, J.; García, B.; Hernández-Rosas, J.I. La planta exótica *Kalanchoe daigremontiana* incrementa el reservorio y flujo de carbono en el suelo. Interciencia 2011, 36, 937–942.

35. Buhle, E.; M, M.; Ruesink, J. Bang for buck: cost-effective control of invasive species with different life histories. Ecological Economics 2005, 52, 355–366.

36. Easterling, M.R.; Ellner, S.P.; Dixon, P.M. Size-specific sensitivity: applying a new structured population model. Ecology 2000, 81, 694–708.

37. Gonzáles, E.; Child, D.; Quintana-Ascencio, P.; Salguero-Gómez, R. Integral projection models. In Demographic Methods across the Tree of life; Salguero-Gómez, R.; Gamelon, M., Eds.; Oxford. University Press: Oxford, 2021; pp. 181–195.

38. Quintana-Ascencio, P.F.; Koontz, S.M.; Smith, S.A.; Sclater, V.L.; David, A.S.; Menges, E.S. Predicting landscape-level distribution and abundance: Integrating demography, fire, elevation and landscape habitat configuration. Journal of Ecology 2018, 106, 2395–2408.

39. Ramula, S.; Rees, M.; Buckley, Y.M. Integral projection models perform better for small demographic data sets than matrix population models: a case study of two perennial herbs. Journal of Applied Ecology 2009, 46, 1048–1053.

40. Quintana-Ascencio, P.; Menges, E.S.; Cook, G.S.; Ehrlé, J.; Afkhami, M. Drivers of demography: past challenges and a promise for a changed future. In Demographic Methods across the Tree of life; Salguero-Gómez, R.; Gamelon, M., Eds.; Oxford University Press: Oxford, 2021; pp. 115–129.

41. Hannan-Jones, M.A.; Playford, J. The biology of Australian Weeds 40. *Bryophyllum Salisb*. species. Plant Protection Quarterly 2002, 17, 42–57.

42. Guillot-Ortiz, D.; Lopez-Pujol, J.; Lumbreras, E.L.; Puche, C. Kalanchoe daigremontiana Raym. Hamet & H. Perrier. ‘Iberian Coast’. Bouteloua 2015, 21, 35–48.

43. García, L.M.; Chávez, L.L. Las Crasuláceas de México; Sociedad Mexicana de Cactología: México D.F. México, 2003.

44. SEMARNAT. Acuerdo por el que se determina la lista de especies exóticas invasoras para México. Diario Oficial de la Federación, 2016. México.

45. Guerra-Garcia, A.; Golubov, J.; Mandujano, M.C. Invasion of *Kalanchoe* by clonal spread. Biological Invasions 2015, 44, 1–8.

46. Capon, R.J.; MacLeod, J.K.; Oelrichs, P.B. Bryotoxins B and C, toxic bufadienolide orthoacetates from flowers of *Bryophyllum tubiflorum* (Crassulaceae). Australian Journal of Chemistry 1995, 39, 1711–1715.

47. MacKenzie, R.A.; Francke, E.P.; Dunster, P.J. The toxicity to cattle and bufadienolide content of six *Bryophyllum* species. Australian Veterinary Journal 1987, 64, 298–301.

48. Williams, M.C.; Smith, M.C. Toxicity of *Kalanchoe* spp. in chicks. American Journal of Veterinary Research 1984, 45, 543–546.

49. Herrera, I.; Hernández, M.J.; Lampo, M.; Nassar, J. Plantlet recruitment is the key demographic transition in invasion by *Kalanchoe daigremontiana*. Population Ecology 2012, 54, 225–237.

50. Williams, J.L.; Crone, E.E. The impact of invasive grasses on the population growth of *Anemone patens*, a long lived native forb. Ecology 2006, 87, 3200–3208.

51. Ren, M.X.; Zhang, D.G.; Zhang, D.Y. Random amplified polymorphic DNA markers reveal low genetic variation and a single dominant genotype in *Eichhornia crassipes* populations throughout China. Weed Research 2005, 45, 236–244.

52. Geng, Y.; Pan, X.; Xu, C.; Zhang, W.; Li, B.; Chen, J.; an Z Song, B.L. Phenotypic plasticity rather than locally adapted ecotypes allow the invasiivse alligator weed to colonize a wide range of habitats. Biological Invasions 2007, 9, 245–256.

53. Lambertini, C.; Riis, T.; Olesen, B.; Clayton, J.S.; Sorrell, B.K.; Brix, H. Genetic diversity in three invasive clonal aquatic species in New Zealand. BMC Genetics 2010, 11, 1–8.

54. Pichancourt, J.P.; van Klinken, R.D. Phenotypic plasticity influences the size, shape and dynamics od the geographic distribution of an invasive plant. PlosOne 2012, 7, e32323.

55. Herrera, I.; Nassar, N. Reproductive and recruitment traits as indicators of the invasive potential of *Kalanchoe daigremontiana* (Crassulaceae) and *Stapelia gigantea* (Apocynaceae) in a Neotropical arid zone. Journal of Arid Environments 2009, 73, 978–986.

56. Schutzenhofer, M.R.; Valone, T.J.; Knight, T.M. Herbivory and population dynamics of invasive and native *Lespedeza*. Oecologia 2009, 161, 57–66.

57. Hahn, M.A.; Buckley, Y.M.; Müller-Scharer, H. Increased population growth rate in invasive polyploid *Centaurea stoebe* in a common garden. Ecology Letters 2012, 15, 947–954.

58. Münzbergová, Z.; Hadincová, V.; Wild, J.; Kindlmannová, J. Variation in the contribution of different life stages to population growth as a key factor in the invasion success of *Pinus strobus*. PLoS ONE 2015, 8, e56953.

59. Tenhumberg, B.; Suwa, T.; Tyre, A.J.; Russell, L.; Louda, S.M. Integral projection models show exotic thistle is more limited than native thistle by ambient competition and herbivory. Ecosphere 2015, 6, 1–18.

60. Van Kleuken, G.; Johnson, S.D. Effects of self-compatiblility on the distribution of invasive European plants in North America. Conservation Biology 2007, 21, 1537–1544.

61. Knight, T.; Havens, K.; Vitt, P. Will the use of less defund cultivars reduce the invasiveness of perennial plants? BioScience 2011, 61, 816–822.

62. González de León, S.; Herrera, I.; Guevara, R. Mating system, population growth, and management for *Kalanchoe pinnata* in an invaded seasonal dry tropical forest. Ecology and Evolution 2016, 6, 4541–4550.

63. Eggli, U. Crassulaceae; Springer-Verlaag: Berlin, Heidelberg, Germany, 2003.

64. Johnson, M.A. The origin of the foliar pseudobulbils in *Kalanchoe daigremontiana*. Journal of The Torrey Botanical Society 1934, 61, 355–366.

65. Guerra-García, A. Evaluación del é clonal en una especie invasora *Kalanchoe delagoensis*. Bachelor Thesis, Facultad de Ciencias, Universidad Nacional Autónoma de México, 2011.

66. Ellner, S.P.; Rees, M. Integral projection models for species with complex demography. The American naturalist 2006, 167, 410–28.

67. R Core Team. R: A Language and Environment for Statistical Computing. R Foundation for Statistical Computing, Vienna, Austria, 2021.

68. Bates, D.; Mächler, M.; Bolker, B.; Walker, S. Fitting Linear Mixed-Effects Models Using lme4. Journal of Statistical Software 2015, 67, 1–48.

69. Stasinopoulos, D.; Rigby, R. Generalized additive models for location scale and shape (GAMLSS) in R. Journal of Statistical Software 2007, 23, 1–46.

70. Jongejans, E.; Jorritsma-Wienk, L.D.; Becker, U.; Dostal, P.; Milden, M.; de Kroon, H. Region versus site variation in the population dynamics of three short-lived perennials. Journal of Ecology 2010, 98, 279–289.

71. Guerra-García, A.; Barrales-Alcalá, D.; Argueta-Guzmán, M.; Cruz, A.; Mandujano, M.C.; Arévalo-Ramírez, J.A.; Milligan, B.G.; Golubov, J. Biomass Allocation, Plantlet Survival, and Chemical Control of the Invasive Chandelier Plant (*Kalanchoe delagoensis*) (Crassulaceae). Invasive Plant Science and Management 2018, 11, 33–39.

72. Griffith, A. Perturbation approaches for integral projection models. Oikos 2017, 126, 1675–1686.

73. Stubben, C.J.; Milligan, B.G. Estimating and analyzing demographic models using the popbio package in R. Journal of Statistical Software 2007, 22, 1e22.

